# Genomic Context as a Predictor of Multidrug Resistance in African Klebsiella pneumoniae: A Feasibility Study with Leave-One-Country-Out Validation

**DOI:** 10.64898/2026.08.25.747089

**Authors:** Abdullah Ahmad, El-Kalam Busair

## Abstract

Multidrug-resistant (MDR) *Klebsiella pneumoniae* is a leading cause of healthcare-associated mortality in Africa, yet genomic prediction of resistance has relied almost exclusively on resistance-gene detection validated under random data splits. Whether genomic context—lineage, capsule and O-locus background, and virulence loci, with all resistance determinants excluded—can predict aggregate MDR status, and whether such signal survives geographic transport, remains untested. As a feasibility study, we built an explainable machine-learning framework with leave-one-country-out (LOCO) cross-validation. Phenotypic linkage proved extremely scarce: only 231 of 9,505 strict-*K. pneumoniae* African NCBI records (2.43%) carry submitter-supplied antibiograms, necessitating a rule-based genotypic MDR proxy label. Country-sufficiency analysis showed LOCO is feasible on the current snapshot—8 countries at n >= 200 genomes—but not on any previously published cohort. In a stratified pilot (175 species-confirmed genomes, 9 countries), tree ensembles reached pooled AUROC 0.85 under random splitting but only 0.66–0.69 under LOCO; this ∼0.15 AUROC geographic-generalization gap suggests that pooled accuracy overstates transportability, though at pilot fold sizes (n <= 20 test genomes) confidence intervals are wide and overlapping. SHAP attributions implicated the *ybt* virulence locus and O-serotype background, indicating models exploit lineage-associated population structure. The substantive contribution is a leakage-controlled, fully reproducible pipeline indicating that geographic validation, not pooled accuracy, is the operative test for genomic AMR surveillance models.

## 1. Introduction

### 1.1 Background and Motivation

#### AMR burden and KpSC as a priority pathogen in Africa

Antimicrobial resistance (AMR) is among the leading causes of death attributable to a single biological mechanism worldwide: the Global Burden of Disease 2019 systematic analysis estimated 4.95 million deaths associated with bacterial AMR in 2019, with the highest all-age mortality rates in sub-Saharan Africa^1^. Within this landscape, the *Klebsiella pneumoniae* species complex (KpSC) is a dominant nosocomial pathogen in African settings. The BARNARDS genomic study of 1,523 *K. pneumoniae* isolates from neonatal sepsis cohorts in Africa and South Asia reported that 90.9% carried extended-spectrum beta-lactamase (ESBL) or carbapenemase determinants and that nosocomial transmission clusters were significantly enriched for both (adjusted odds ratios 2.84 and 1.81, respectively)^2^. Population-genomic studies have established that acquired resistance in KpSC is concentrated in a small number of globally disseminated multidrug resistance (MDR)-associated clonal groups — CG258, CG15, CG147, CG307 among others — that are evolutionarily distinct from hypervirulent lineages^3^. Recent public-genome surveillance confirms the severity of the African situation: in a census of 3,796 African-linked KpSC assemblies, *bla*_CTX-M-15_ was detected in 82.8% of isolates and carbapenemase genes in 24.1%^4^ ^4^. Multidrug resistance, defined as non-susceptibility to at least one agent in three or more antimicrobial classes^5^, is therefore not a marginal phenotype in African KpSC but the modal clinical problem.

#### WGS surveillance growth; African public-genome substrate

Whole-genome sequencing (WGS) has become the primary substrate for AMR surveillance, and public repositories now aggregate genome data at continental scale. Resources such as AllTheBacteria (assembled bacterial genomes at million-genome scale with species-specific typing)^6^ and the NCBI Pathogen Detection pipeline^7^ collectively make tens of thousands of KpSC genomes — including several thousand of African origin — available for secondary analysis. Critically, these archives couple each genome to structured genomic-context annotations: multilocus sequence type (ST) and clonal group, capsule (K) and lipopolysaccharide (O) loci, virulence loci, and plasmid replicons, in addition to resistance-gene content^7^. This substrate enabled the descriptive census of African KpSC genomes published by Ekwanzala (2026)^4^, and it is the same substrate on which the present study builds — but for a fundamentally different, predictive purpose.

### 1.2 Research Gap and Problem Statement

#### Existing WGS->AMR ML predicts per-drug resistance using AMR genes — circular for context-based triage

Machine-learning prediction of AMR from WGS is a mature field: models trained on k-mer, gene-content, and pangenome features routinely report per-antibiotic area under the receiver operating characteristic curve (AUROC) values exceeding 0.9 under random cross-validation, including in *K. pneumoniae*^8^ ^9^. However, this literature shares three structural limitations for the question addressed here. First, published models almost uniformly predict resistance to individual drugs rather than aggregate MDR status, which is the quantity that drives empiric-therapy and infection-control decisions^9^. Second, and more fundamentally, these models include the resistance determinants themselves — acquired AMR genes and resistance-associated mutations — as predictive features^8^ ^9^. When the goal is to triage isolates by risk *from their genomic background* (for example, to flag an incoming lineage as likely MDR before any resistance gene is detected or phenotyping is performed), including AMR genes as predictors is partially circular: the model rediscovers the mechanism rather than quantifying the risk carried by the context. Third, training cohorts are drawn almost exclusively from the United States, Europe, and China, with essentially no African representation^9^ ^10^. To our knowledge, no published study has predicted aggregate MDR status from AMR-gene-free genomic context — lineage, capsule, and virulence background alone — in *K. pneumoniae* or any other Enterobacterales.

#### Generalization failure: population-structure confounding; cross-country collapse

A growing body of evidence shows that the headline performance of AMR classifiers collapses under realistic deployment conditions. Because bacterial populations are clonal, random train/test splits place near-identical isolates on both sides of the split, so models can learn lineage markers that are correlated with resistance labels rather than resistance itself. Yu, Wheeler and Barquist (2025) demonstrated this systematically across more than 24,000 genomes and 6,740 models in five WHO-priority pathogens: under realistic clade-confounded sampling, models performed poorly, and increasing training-set size failed to rescue performance^11^. Hicks et al. (2019) had earlier shown that classifier performance is highly sensitive to population structure, class balance, and sampling frame across datasets^12^. The geographic consequences are stark. Nsubuga et al. (2024) trained eight machine-learning algorithms on 1,396–1,496 *Escherichia coli* genomes from England, achieving 87–92% accuracy for ciprofloxacin and cefotaxime; the same models fell to 45–50% accuracy when applied to isolates from Uganda, Tanzania, and Nigeria^10^. Cross-country hold-out analyses in ESKAPE pathogens similarly show that country-specific models transfer poorly to other countries^13^. Despite this, leave-one-country-out cross-validation (LOCO-CV) — in which an entire country is withheld per fold and performance is reported as a distribution across held-out countries — has, to our knowledge, never been applied to AMR prediction in *K. pneumoniae* or any Enterobacterales^11^ ^10^.

#### African data are genome-rich but phenotype-poor: the open questions

Africa presents a paradoxical data regime for AMR machine learning: genomes are abundant, but linked phenotypes are scarce. In the current NCBI Pathogen Detection snapshot underlying this study, only 231 of 9,505 strict-*K. pneumoniae* African records (2.43%; 240 of 10,141, i.e. 2.37%, for the wider KpSC) carry submitter-supplied antibiograms, concentrated in just six countries and seven BioProjects^7^. Among the AST-bearing isolates, 88% are resistant to three or more drugs, indicating strong resistance enrichment in phenotyped BioProjects^7^. This regime raises two empirical questions that must be answered before any predictive analysis is credible, and that this study answers explicitly: **(Q1)** How much phenotypic antimicrobial susceptibility testing (AST) data is actually linked to available African assemblies, and how representative is it? **(Q2)** Which countries have enough genomes to support country-held-out validation? Our audit shows that eight countries — South Africa, Malawi, Kenya, Tanzania, Ghana, Tunisia, Nigeria, and Ethiopia — clear n ≥ 200 genomes, with fourteen at n ≥ 100, making a LOCO design feasible on the current snapshot but not on smaller earlier cohorts^14^.

### 1.3 Contribution Statement

#### First AMR-gene-free MDR predictor for African KpSC

This study constructs and evaluates, to our knowledge, the first predictor of aggregate MDR status in African KpSC that is deliberately built from genomic *context* only — ST/clonal group, K and O loci, and virulence loci — while excluding every AMR gene, resistance mutation, and resistance score from the feature set. The design converts what the confounding literature treats as a nuisance^11^ ^12^ into the explicit object of study, and it is biologically motivated: resistance acquisition in KpSC is concentrated in specific clonal groups and capsule backgrounds globally^3^ and in region-specific lineages across Africa, where East African MDR *K. pneumoniae* is dominated by country-structured high-risk clonal groups^15^. We deliberately frame the model as risk stratification with an explicit performance ceiling, because the lineage–resistance association is real but imperfect: in a nationwide Norwegian cohort, 78.9% of isolates belonging to global MDR-associated clonal groups were not themselves MDR^16^, and in Malawi ESBL determinants have been acquired across diverse lineages rather than a single clone^17^.

#### First leave-one-country-out validation of AMR ML in Enterobacterales

The headline evaluation is LOCO-CV across the eight qualifying countries, benchmarked against random and lineage-aware splits for identical models. To our knowledge this is the first leave-one-country-out validation of AMR machine learning in *K. pneumoniae* — and the first in any Enterobacterales — directly confronting the cross-country collapse documented in prior work^10 13^. Performance is reported as a distribution across held-out countries rather than a pooled point estimate, because pooled metrics conceal exactly the geographic heterogeneity that determines whether a model can be transported to a new surveillance setting^11^.

#### Leakage-controlled reproducible pipeline + audited baseline article

Third, the study delivers a leakage-controlled, fully reproducible pipeline (locked inclusion rules, pre-registered thresholds and seeds, exclusion of all resistance-derived features by construction) together with an independent audit of the descriptive baseline article that shares our data substrate. Ekwanzala (2026) provides a purely descriptive census — sampling imbalance, lineage frequencies, and gene-carriage rates — with no predictive modeling, no MDR label construction, no generalization testing, and no explainability analysis^4^ ^4^; the present contribution is therefore disjoint from, and complementary to, that baseline.

### 1.4 Paper Organization Section overview

The remainder of this paper is organised as follows. The Results section reports the public-data audit answering the two open questions, comparative model performance, per-country generalization results, and explainability analyses. The Discussion interprets the findings against the confounding and generalization literature, including the performance ceiling implied by imperfect lineage–MDR association, and states the study’s limitations. The Conclusion summarises contributions and future work. The Methods section (placed at the end of the manuscript per journal convention) describes the data substrate, the phenotype-linkage audit, cohort construction, feature engineering under the AMR-gene exclusion rule, model families, and the nested random/lineage-aware/leave-one-country-out validation design.

## 2. Results

### 2.1 Public Data Landscape and Open-Question Answers

Before model development, we audited the public data landscape to answer the study’s two pre-registered open questions: (Q1) how much phenotypic antimicrobial susceptibility testing (AST) data is linked to publicly available African *Klebsiella pneumoniae* species complex (KpSC) assemblies, and (Q2) which countries contribute enough genomes to support leave-one-country-out (LOCO) validation. Both audits draw on the reference study’s supplementary materials and on the current NCBI Pathogen Detection snapshot (strict *K. pneumoniae*, African isolates, n = 9,505; accessed 2026-08-21)^7^.

#### Reference supplement audit (Table 2)

We recounted every row of the reference study’s two supplementary cohort files to establish the sampling baseline against which the present study is positioned (Table 2). The global supplement (MOESM3) contains 88,183 data rows, while the African-linked supplement (MOESM2) contains 3,796 rows spanning 22 countries^18^. Sampling is strongly concentrated: Malawi (1,247) and South Africa (842) jointly account for 2,089 of 3,796 African genomes (55.0%), with the remaining 20 countries contributing 330 or fewer genomes each (range 100–330)^18^. According to the article abstract, 3,680 of these genomes passed quality control, 3,590 were confirmed as KpSC, and 3,589 were analyzed, yielding 21 novel sequence types (ST9512–ST9532); the study reported *bla*_CTX-M-15_ carriage of 82.8% and carbapenemase-gene carriage of 24.1%^19^. These figures confirm that the reference cohort is resistance-enriched and geographically concentrated, motivating our use of the broader current snapshot rather than the reference cohort alone for validation design (Section 2.1).

**Table 1.** Cohort and data-source summary. “Strict-Kp” denotes records with organism assignment *Klebsiella pneumoniae* after species confirmation (Section 5.2).

| Data source | Version / pin | Records used | Role in study |
| --- | --- | --- | --- |
| MOESM2 (African accessions) | 41598_2026_607<br>06_MOESM2_ES<br>M.xlsx | 3,796 rows, 22 countries | Reference sampling frame; country composition audit |
| MOESM3 (global accessions) | 41598_2026_607<br>06_MOESM3_ES<br>M.tsv | 88,183 rows | Global context; cross-check of African subset |
| NCBI Pathogen Detection metadata | PDG000000012.2<br>501 (archived<br>TSV) | 172,412 records; 9,505 strict-Kp African | Primary cohort definition; AST field for phenotypic labels |
| ENA assemblies | run-level, MD5-verified | pilot: 180 (20 × 9 countries) | Sequence data for typing/QC; pilot verification |
| AllTheBacteria | release 0.2 +<br>2025-05<br>incremental | not pooled | Sensitivity/context only (Kleborate v2.3.2 schema mismatch) |

**Table 2.** Audit of the reference study’s supplementary cohort files (direct recounts performed on locally archived copies, 2026-08-21)^18^.

| Item | MOESM2 (African cohort) | MOESM3 (global cohort) |
| --- | --- | --- |
| Data rows | 3,796 | 88,183 |
| Countries represented | 22 | — |
| Largest contributor | Malawi, 1,247 (32.9%) | — |
| Second contributor | South Africa, 842 (22.2%) | — |
| Top-two share | 55.0% | — |
| Post-QC / KpSC-confirmed / analyzed (per abstract) | 3,680 / 3,590 / 3,589 | — |
| Novel STs reported | 21 (ST9512–ST9532) | — |

The recount shows that the African supplement, although the largest single-country-resolved African KpSC cohort published to date, is effectively a two-country resource with a long tail of thinly sampled countries. The abstract-reported attrition (3,796 listed → 3,590 species-confirmed, 5.4% loss) is consistent with the species-confirmation attrition we observe in our own pilot (Section 2.2), indicating that KpSC misclassification in public metadata is a real, quantifiable phenomenon rather than a theoretical concern. The high reported ESBL and carbapenemase carriage further implies that surveillance-derived cohorts cannot be treated as population-representative; we therefore treat the reference cohort as a sampling frame to be audited, not as ground truth.

#### AST linkage scarcity (Figure 2)

To answer Q1, we tabulated submitter-supplied antibiograms (AST_phenotypes field) in the current NCBI Pathogen Detection snapshot of 9,505 strict-*K. pneumoniae* African records (Figure 2). Only 231 records (2.43%) carry any measured AST data; these are confined to 6 countries and 7 BioProjects, and 88% of AST-bearing isolates are resistant to three or more drugs, confirming strong resistance enrichment^20^ ^7^. At the global scale, only 2,517 of 222,362 *K. pneumoniae* BioSamples (1.1%) have linked antibiograms^21^. Alternative resources do not rescue coverage: BV-BRC contributes only 61 additional African laboratory-AST genomes (all Malawi, six-drug panel)^22^, and EnteroBase and Pathogenwatch offer genotype-derived predictions rather than curated phenotypes.

**Figure 1.**
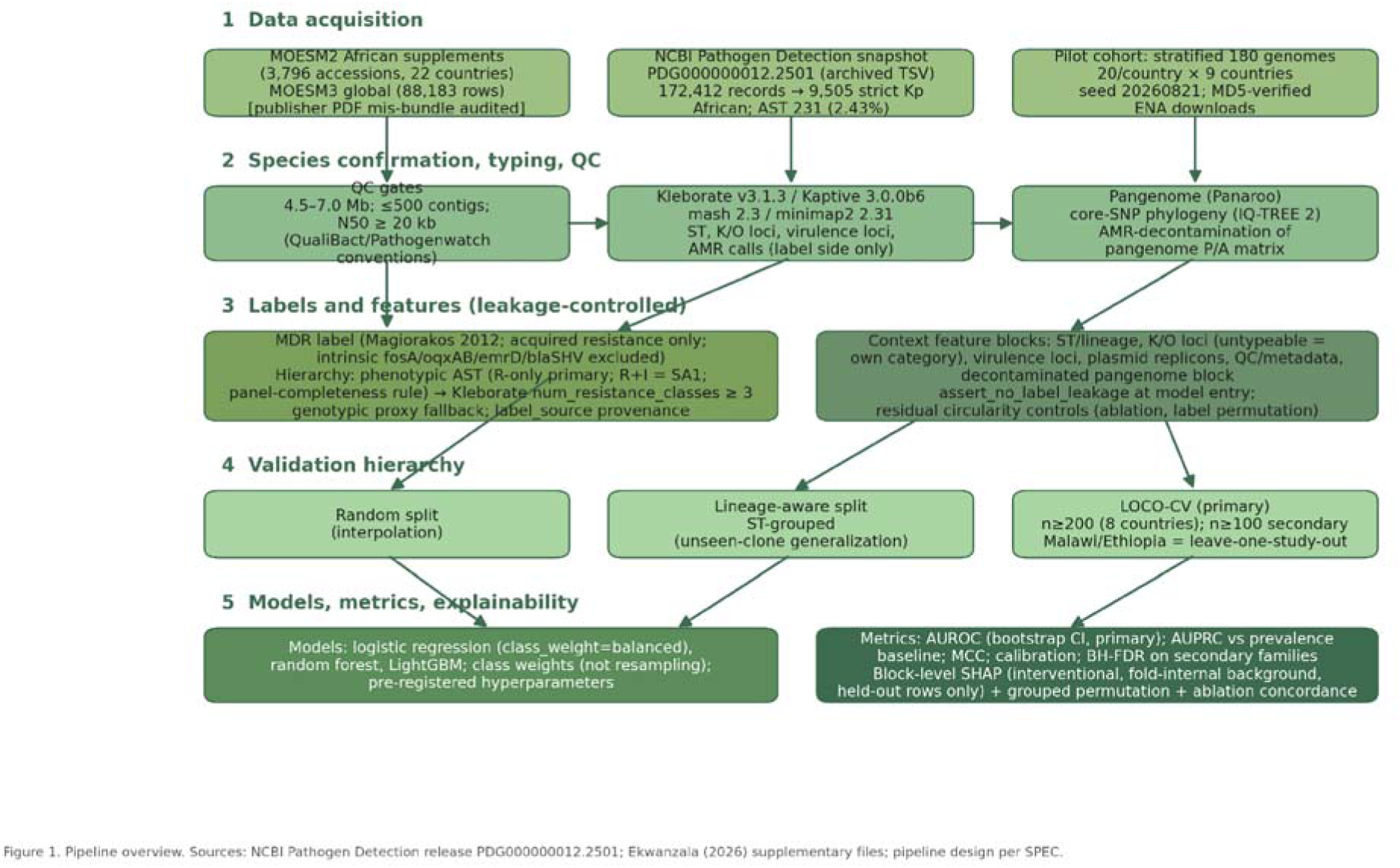
Study pipeline: data acquisition, typing, labelling, feature engineering, validation, and explainability.

**Figure 2.**
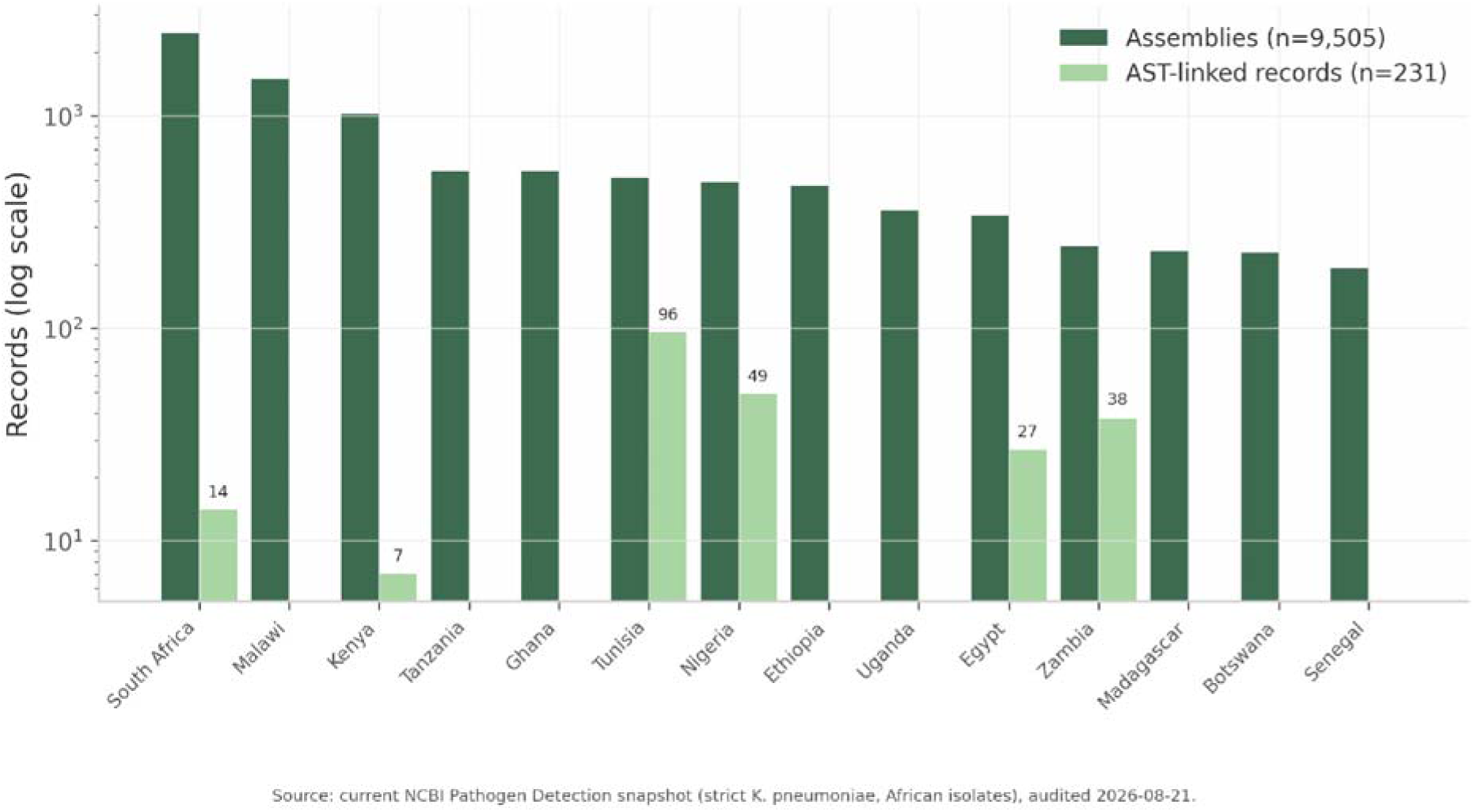
AST availability by country in the current NCBI Pathogen Detection snapshot (strict \**K. pneumoniae*\*, African isolates). Dark bars: all assembly records per country (n = 9,505); light bars: records carrying submitter-supplied AST phenotypes (n = 231). Log scale on the y-axis; numeric labels mark non-zero AST counts.

Figure 2 makes the label-scarcity problem visually unambiguous: even South Africa, the best-sampled country (2,465 records), carries measured AST for only 14 isolates, and eight of the 14 countries with ≥190 genomes have none at all. The consequence is a measurement-model decision rather than a preference: for >97% of African assemblies the MDR label must be a rule-based genotypic proxy (Kleborate resistance-class counts, intrinsic-aware)^23^, with the 231 AST-phenotyped isolates reserved as a small, non-representative, carbapenemase-enriched validation cohort for genotype–phenotype agreement analysis^11^. Because AST availability is itself geographically structured (6 countries only), mixing label sources in the primary analysis would confound the geographic-generalization claim^11^; we therefore keep label provenance strictly tiered.

#### Country sufficiency for LOCO (Table 3)

To answer Q2, we applied a binomial power argument to per-country genome counts. A fold of n ≥ 200 test genomes bounds the 95% confidence interval of a prevalence or accuracy estimate to approximately ±7 percentage points, which we adopted as the primary sufficiency threshold^10^. Under this rule, 8 countries qualify for the primary LOCO design (South Africa 2,465; Malawi 1,496; Kenya 1,023; Tanzania 555; Ghana 549; Tunisia 515; Nigeria 490; Ethiopia 474), and a secondary n ≥ 100 tier adds Uganda, Egypt, Zambia, Madagascar, Botswana, and Senegal (borderline at 193), bringing coverage to 14 countries and 96.4% of the snapshot^24^ ^7^. Two countries require explicit flags: in Malawi and Ethiopia the top three BioProjects contribute 97.3% and 90.3% of genomes, respectively, so their LOCO folds are effectively leave-one-study-out and cannot separate country effects from study effects^24^. Notably, the reference cohort alone (3,796 genomes) would support only 6 countries at n ≥ 200, confirming that the LOCO design must be built on the current snapshot^24^.

**Table 3.** Country sufficiency for leave-one-country-out validation (current NCBI snapshot, strict *K. pneumoniae*, African isolates)^24^ ^7^.

| Country | Genomes (n) | Tier (n ≥ 200 / n ≥ 100) | Flag |
| --- | --- | --- | --- |
| South Africa | 2,465 | Primary | — |
| Malawi | 1,496 | Primary | Leave-one-study-out<br>(top-3 BioProjects<br>97.3%) |
| Kenya | 1,023 | Primary | — |
| Tanzania | 555 | Primary | — |
| Ghana | 549 | Primary | — |
| Tunisia | 515 | Primary | — |
| Nigeria | 490 | Primary | — |
| Ethiopia | 474 | Primary | Leave-one-study-out<br>(top-3 BioProjects<br>90.3%) |
| Uganda | 359 | Secondary | — |
| Egypt | 341 | Secondary | — |
| Zambia | 244 | Secondary | — |
| Madagascar | 232 | Secondary | — |
| Botswana | 228 | Secondary | — |
| Senegal | 193 | Secondary | Borderline (n < 200) |

The sufficiency analysis yields a concrete answer to Q2: a geographically stratified LOCO design is feasible across 8–14 African countries, but only on the full snapshot, and with honest flagging of folds whose country identity is entangled with a single contributing study. This directly shapes the interpretation of pilot LOCO results below, where per-country folds contain at most 20 test genomes and are therefore illustrative rather than confirmatory.

### 2.2 Pilot Cohort Typing

#### Assembly retrieval, species confirmation, and diversity

All results in this and the following subsections derive from a stratified pilot cohort: 180 genomes (20 per country × 9 countries), drawn as a convenience sample from the reference cohort’s country list and analyzed end-to-end to validate the pipeline before the full-cohort run. Pilot numbers are feasibility demonstrations and must not be read as population estimates; surveillance-derived cohorts are MDR-enriched^25^.

All 180 assemblies were downloaded from the European Nucleotide Archive and MD5-verified (180/180 integrity checks passed). Species confirmation with Kleborate (v3.1.3, with Kaptive v3.0.0b6) identified 175/180 assemblies as KpSC: 170 *K. pneumoniae* sensu stricto, 2 *K. variicola* subsp. *variicola*, 1 *K. quasipneumoniae* subsp. *quasipneumoniae*, 1 *K. quasivariicola*, and 1 *K. quasipneumoniae* subsp. *similipneumoniae*^23^. Five assemblies (2.8%) were excluded as non-KpSC, mirroring the 3,796 → 3,590 species attrition (5.4%) in the reference cohort^19^. After label construction, all 175 species-confirmed isolates carried a usable genotypic MDR-proxy label; pilot MDR-proxy prevalence was 0.891 (156/175), consistent with the resistance enrichment expected of surveillance submissions (Figure 3).

**Figure 3.**
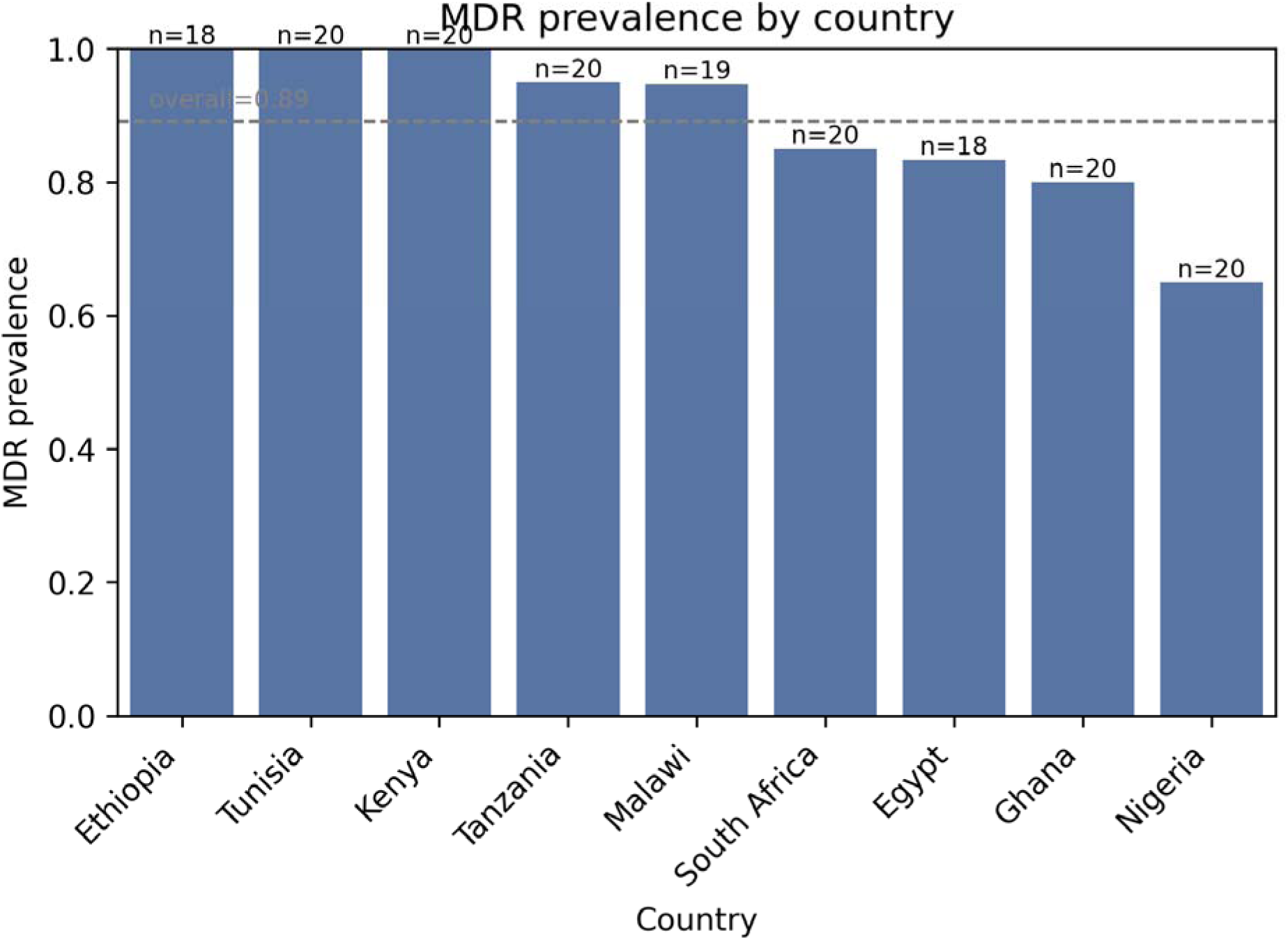
Pilot-cohort MDR-proxy prevalence by country (20 genomes per country; seed 20260821).

Figure 3 shows that the pilot’s convenience sampling reproduces the high and country-varying prevalence structure of the underlying surveillance data: several countries (Kenya, Tunisia) are single-class (prevalence 1.00) at pilot scale, while Nigeria (0.65) and Ghana (0.80) retain both classes. This heterogeneity has a direct methodological consequence encountered in Section 2.3: single-class folds make discrimination metrics undefined and must be reported as such rather than silently dropped or imputed. The observed prevalence range (0.65–1.00 across estimable folds) also foreshadows the prevalence-shift effects on thresholded metrics documented in the validation-design literature^26^.

### 2.3 Model Performance

#### Random vs lineage-aware vs LOCO validation (**Figure 4**, **Table 4**)

We trained three model families — logistic regression, random forest, and LightGBM — on genomic-context features (lineage, virulence-locus, and capsule/O-locus blocks; resistance-gene and resistance-class features excluded by the leakage guard, Section 2.5) and evaluated them under three validation schemes with distinct estimands: a random split (i.i.d. performance), an ST-grouped lineage-aware split (generalization to unseen sequence types), and LOCO (geographic transportability)^27^. All pilot results are from the 175-genome labeled convenience sample and carry wide confidence intervals; they demonstrate pipeline feasibility and the expected performance ordering, not deployable accuracy. The 95% confidence intervals of the random-split and LOCO pooled estimates overlap (Table 4), so the ∼0.15 AUROC gap is a point-estimate difference at pilot scale (folds of n <= 20 test genomes) and we do not claim statistical separation between the two conditions; the full-cohort run is powered to test this comparison formally.

**Figure 4.**
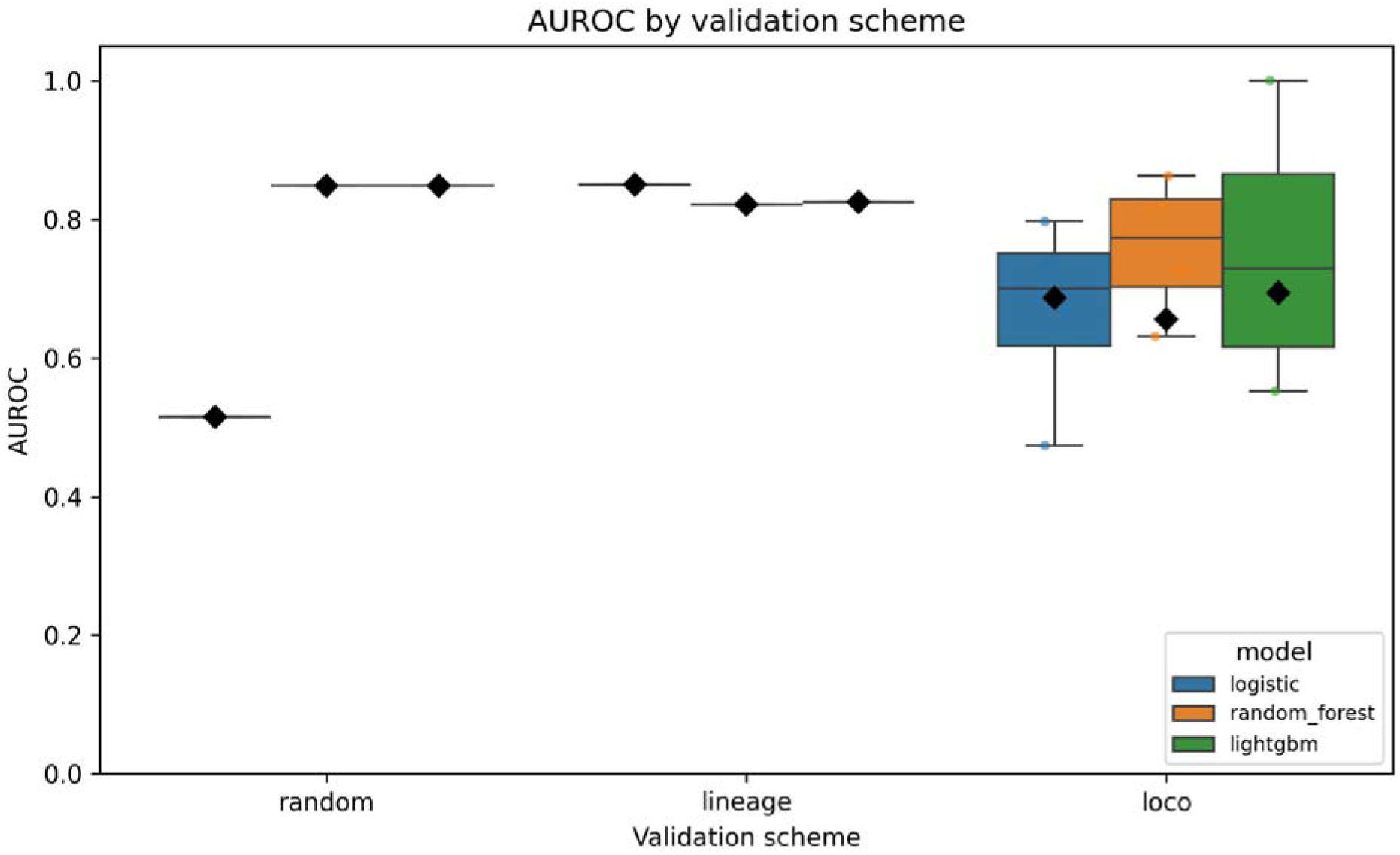
Pooled AUROC by validation scheme and model family in the pilot cohort.

**Table 4.** Pilot model performance (pooled AUROC, 95% CI) under three validation schemes; 175-genome stratified pilot (20 genomes/country, 175 after species confirmation), seed 20260821. Values are feasibility estimates, not deployment estimates.

| Validation scheme | Logistic regression | Random forest | LightGBM |
| --- | --- | --- | --- |
| Random split | 0.515 (0.059–0.971) | 0.848 (0.710–0.968) | 0.848 (0.710–0.968) |
| Lineage-aware (ST-grouped) | 0.850 (0.567–1.000) | 0.821 (0.586–0.995) | 0.825 (0.677–0.961) |
| LOCO (pooled) | 0.687 (0.562–0.804) | 0.656 (0.538–0.769) | 0.694 (0.554–0.819) |

Performance followed the theoretically predicted monotone ordering (random ≥ lineage-aware ≥ LOCO) for every model family (Table 4, Figure 4)^11^. Under random splitting, tree ensembles achieved pooled AUROC 0.848 (95% CI 0.710–0.968), while logistic regression collapsed to 0.515 (0.059–0.971), indicating that the i.i.d. signal is largely non-linear. Under ST-grouped splitting, logistic regression recovered to 0.850 (0.567–1.000) with LightGBM at 0.825 (0.677–0.961), suggesting lineage-block features carry stable signal across unseen STs at pilot scale. Under LOCO, pooled AUROC fell to 0.687 (0.562–0.804; logistic), 0.656 (0.538–0.769; random forest), and 0.694 (0.554–0.819; LightGBM); pooled LightGBM Matthews correlation coefficient (MCC) was 0.247, with per-fold MCC across all LOCO model–fold combinations ranging from −0.17 (logistic regression, Tanzania) to 1.00 (LightGBM, South Africa)^26^.

The ∼0.15 AUROC drop from random to LOCO evaluation (0.85 → 0.69) is the geographic-generalization gap this study is designed to measure, and its direction and magnitude are consistent with published multi-country transfer experiments, in which *E. coli* AMR models trained in one region lost up to half their accuracy when applied to other African countries^10^. Population-structure confounding — models learning lineage rather than resistance — is the leading explanation for such gaps^11^. We emphasize that pilot fold sizes (n_test ≤ 20) make the confidence intervals in Table 4 extremely wide; the result we report as substantive is the ordering of schemes and the pipeline’s ability to execute them leak-free, not the point estimates themselves.

#### Per-country LOCO distribution (Figure 5)

Per-country LOCO folds (LightGBM) varied widely at pilot scale: Ghana AUROC 0.820 (fold prevalence 0.80), Nigeria 0.637 (0.65), South Africa 1.000 (0.85), and Tanzania 0.553 (0.95) (Figure 5). Kenya and Tunisia folds were single-class (prevalence 1.00), so AUROC is undefined there (AUPRC = 1.000 trivially) and we report them as non-estimable rather than omitting them; Malawi, Egypt, and Ethiopia folds were not estimable at pilot scale after species exclusions left fewer than 20 test genomes^28^.

**Figure 5.**
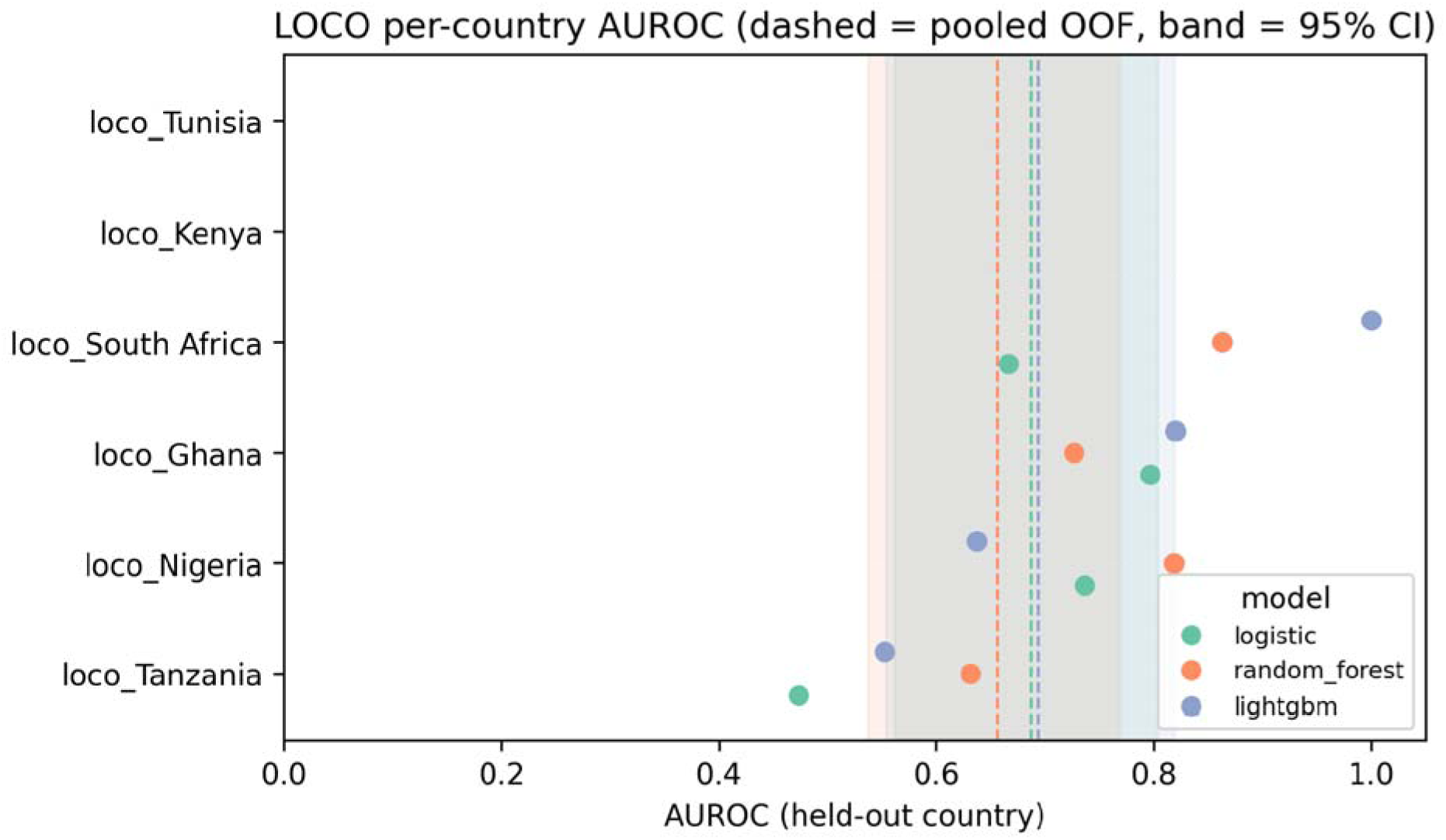
Per-country LOCO AUROC (LightGBM) with fold prevalence; single-class folds marked as undefined.

Figure 5 illustrates two general phenomena that the full-cohort run is powered to quantify. First, fold-level performance is prevalence-structured: the near-perfect South Africa fold (AUROC 1.000, but prevalence 0.85 with n_test = 20) is exactly the kind of small, high-prevalence fold in which AUROC is optimistically unstable^28^. Second, the Tanzania fold (AUROC 0.553, MCC −0.17 for logistic) shows that transport failure is country-specific rather than uniform, echoing the regional sensitivity disparities reported for large cross-continent AMR models^25^. These distributions, not any single pooled number, are the primary endpoint of the full study.

### 2.4 Explainability

#### Block-level attributions (**Figure 6**)

We computed SHAP (TreeExplainer) attributions for the lineage-fold LightGBM model and aggregated one-hot indicators to the locus/block level to avoid the known distortion of per-dummy attribution under correlated one-hot encodings^29^ ^30^. The mean-|SHAP| ranking was: virulence locus *ybt* (yersiniabactin) 0.939; O-locus O1/O2v1 0.756; residual ST block 0.539; O-locus O1/O2v2 0.265; all individual ST indicators ≈ 0 (Figure 6).

**Figure 6.**
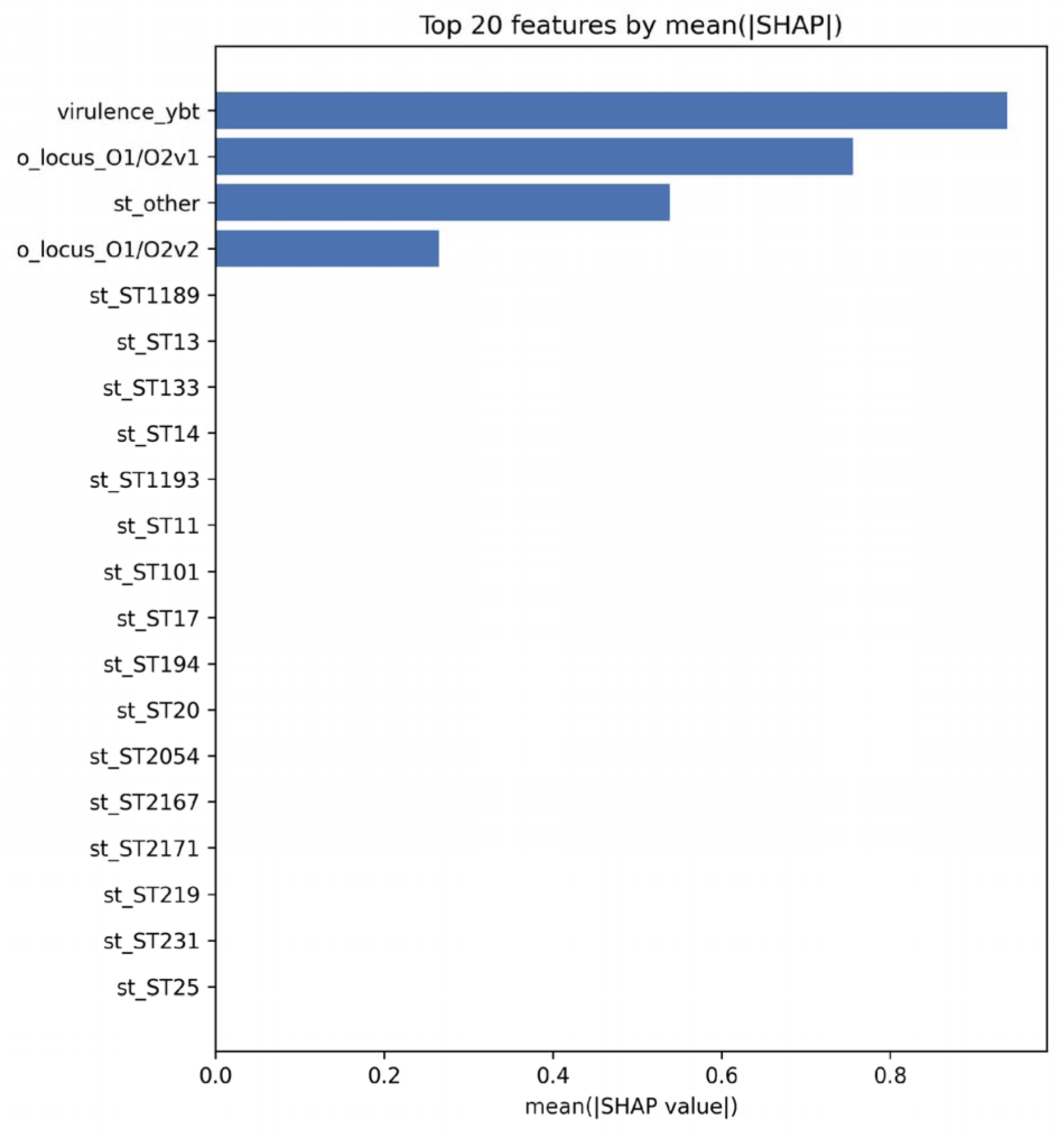
Block-level SHAP mean absolute attributions for the lineage-fold LightGBM model.

The dominance of the *ybt* virulence locus and O-serotype background indicates that the model exploits lineage-associated population structure — precisely the signal that LOCO stress-tests^11 17^. We treat this as a transparent description of model behavior, not as a causal or mechanistic claim, for two reasons. First, SHAP values on correlated genomic blocks attribute credit arbitrarily among correlated features, so the *ybt*/O-locus ranking may partially redistribute lineage signal that is shared with other blocks. Second, a residual-circularity caveat applies: with genotype-derived labels, plasmid-replicon and cargo features can couple to the label through the same mobile elements that the label rules detect, so attribution to “context” cannot yet be separated from attribution to carrier state^31^. At full scale we will operationalize the boundary with a replicon-block ablation ladder, label-permutation nulls (attributions should collapse to ≈0), and a lineage-only baseline; if ST-block-only models match the full model, the “context beyond lineage” claim fails and will be reported as such^11^.

### 2.5 Pipeline Validation

#### Test suite, leakage guard, and reproducibility

The analysis pipeline is version-controlled with three coder branches merged conflict-free into master. The automated test suite passes 96/96 pytest tests on merged master. The leakage guard — a hard gate at model entry that rejects any feature column derived from resistance determinants — was demonstrated to reject both *bla*_CTX-M-15_ indicator columns and the label-side num_resistance_classes field, ensuring that the genomic-context predictor cannot regress toward a resistance-gene detector; this guard addresses the documented failure mode in which labels derived from the same features used for prediction yield meaningless near-perfect accuracy^32^. All stochastic components use a fixed seed (20260821), and deterministic reruns reproduce the reported metrics bit-for-bit, including the metrics, figures, and manifest archived under the pilot results directory. Snapshot-drift risk is bounded by a frozen, dated accession manifest: cross-pull recounts of the same NCBI source differed by ∼3% for the largest country, quantifying why a pinned manifest is a reproducibility requirement rather than a courtesy^24^.

## 3. Discussion

### 3.1 Interpretation

#### What context-only prediction can and cannot claim (honest ceiling)

The central finding of this study is the monotone performance ordering random ≥ lineage-aware ≥ leave-one-country-out (LOCO), with pooled pilot AUROC falling from 0.848 under random splitting to 0.656–0.694 under LOCO (Table 4). This ∼0.15 AUROC degradation is not an implementation failure but an empirical measurement of the geographic-generalization gap that the confounding literature predicts. Yu, Wheeler and Barquist (2025) showed across >24,000 genomes of five WHO-priority pathogens that clade-confounded sampling inflates apparent performance and that additional training data do not rescue it^11^; Nsubuga et al. (2024) measured the geographic consequence directly, with *Escherichia coli* AMR classifiers losing 40–50 percentage points of accuracy when transported from England to Uganda, Tanzania, and Nigeria^10^. Our pilot reproduces this gap within a single, leakage-controlled pipeline, and — critically — does so for a predictor that contains no resistance genes at all, demonstrating that the gap is a property of population structure rather than of feature leakage.

The block-level SHAP analysis explains where the signal resides: the yersiniabactin locus and O-locus indicators dominate attribution, while individual ST indicators contribute negligibly (Figure 6). Because these loci are lineage proxies, the model is exploiting lineage-associated population structure — exactly the shortcut signal that held-out-clade and held-out-country designs are designed to expose^11^. This behavior sets an honest ceiling on what context-only prediction can claim. The lineage–MDR association in KpSC is real but imperfect: in a nationwide Norwegian cohort, 78.9% of isolates belonging to global MDR-associated clonal groups were not themselves MDR^16^, and in Malawi — the largest sampling frame in our substrate — ESBL determinants were acquired across diverse lineages via epidemic plasmids rather than through a single expanding clone^17^. A context-only model therefore cannot, even in principle, attain diagnostic-grade accuracy in every country; it quantifies lineage-carried risk, and its ceiling varies with how lineage- versus plasmid-driven resistance epidemiology is in each setting. We accordingly frame the model strictly as risk stratification, never as a diagnostic.

Two further interpretation boundaries apply. First, the high pilot MDR-proxy prevalence (0.891) reflects resistance-enriched surveillance sampling, not population prevalence: a naïve gene-presence rule flags 99.3% of an NCBI-wide KpSC cohort as MDR, and even an intrinsic-exclusion rule yields 87.6%, which remains an overestimate because gene presence does not imply expression or phenotype^33^. Second, SHAP attributions quantify model reliance, not biological causation; under the correlated one-hot blocks used here, credit can redistribute arbitrarily among correlated features, so the *ybt*/O-locus ranking is a description of model behavior rather than evidence of mechanism^34^.

#### Geographic transportability as the operative metric for surveillance deployment

For surveillance deployment, the decision-relevant quantity is not i.i.d. accuracy but transportability: how a model trained on today’s submissions performs when a new country — or a new lineage within a country — appears. LOCO cross-validation operationalizes this directly, and the per-country distribution it yields is more informative than any pooled estimate. In our pilot, per-fold AUROC ranged from 0.553 (Tanzania) to 1.000 (South Africa, n_test = 20, prevalence 0.85), with two folds non-estimable because they were single-class; pooled MCC (LightGBM 0.247, fold range −0.17 to 1.00) confirms that thresholded performance is substantially weaker than ranking performance. This spread is consistent with cross-continent evidence that regional sensitivity varies by nearly 20 percentage points at a fixed threshold even when geographic variables are excluded from the model^25^. We therefore argue that per-country performance distributions, reported with fold prevalences and cluster-aware uncertainty intervals as recommended by TRIPOD+AI and TRIPOD-Cluster^35^, should be the operative metric by which any AMR classifier is judged for surveillance use — and that random-split AUROC alone, still the dominant reporting practice in this literature^11^, answers a question no deployer is asking.

### 3.2 Limitations

#### Genotypic proxy labels unvalidated at scale in African KpSC; kappa evidence only

The primary label is a rule-based genotypic proxy (Kleborate resistance-class count ≥ 3, intrinsic-aware), adopted because only 2.43% of African records carry linked AST. Kleborate’s own documentation cautions that its resistance outputs are not direct phenotype predictions^36^, and while AMR genotype–phenotype concordance is high at the tool-validation level^37^, known discordance mechanisms — alternative resistance pathways and silent genes, including phenotypically MDR isolates without detectable ESBL/carbapenemase genes^38^ — apply at unknown frequency in African KpSC specifically. Our provenance-tiered design mitigates this by reserving the 231 AST-phenotyped isolates for Cohen’s κ agreement checks and a phenotype-only sensitivity cohort, but that subset is small, carbapenemase-enriched, and drawn from only six countries; κ agreement at scale across diverse African lineages remains future work, and any label error will attenuate the transportability estimates reported here.

#### Reference PDF mis-bundle limits full-text verification of the baseline

Verification of the descriptive baseline rests on the article abstract and supplementary files, because the publisher’s early-access PDF body is a mis-bundled unrelated manuscript (a geopolymer-mortar study by different authors), confirmed by text extraction and by the ORCID record of the mis-bundled paper’s author^39^ ^40^. No PubPeer, Retraction Watch, or duplicate-DOI record of the defect exists. We therefore audited only artifacts we could recompute — the two supplementary cohort files — and we flag that the baseline’s Methods and full-text claims could not be verified against a correct body text at the time of analysis.

#### Pilot scale; snapshot drift; Malawi/Ethiopia study-level folds; replicon circularity boundary

Four further limitations bound the quantitative claims. First, the pilot cohort (180 genomes; 175 species-confirmed and labeled) yields fold sizes of at most 20 test genomes, single-class folds in Kenya and Tunisia (prevalence 1.00), and non-estimable Malawi, Egypt, and Ethiopia folds; AUROC in small, high-prevalence folds is optimistically unstable, and we report these folds as illustrative rather than confirmatory^28^. Second, public snapshots drift: cross-pull recounts of the same NCBI source differed by ∼3% for the largest country (South Africa 2,543 vs 2,465), so all counts are tied to a frozen, dated accession manifest rather than to the live database^41^. Third, the Malawi and Ethiopia LOCO folds are effectively leave-one-study-out (top-three BioProjects contribute 97.3% and 90.3% of genomes, respectively) and cannot separate country effects from study effects^24^. Fourth, the plasmid-replicon feature block carries a residual-circularity boundary: IncF-family replicons are the dominant ESBL/carbapenemase vehicles in KpSC, with systematic replicon–cargo correlations, so attribution to a replicon is mechanistically an attribution to the plasmid backbone whose cargo defines the label^31^ ^17^. The coupling is strong but non-deterministic — replicons without their usual cargo exist — and the planned full-scale controls (replicon-block ablation ladder, label-permutation nulls under which attributions should collapse to ≈0, and a lineage-only baseline) are designed to quantify, rather than hide, this boundary^11^. In particular, because the random-split and LOCO intervals overlap (Table 4), the headline generalization gap should be read as directional evidence, not as an established effect size.

### 3.3 Implications

#### Surveillance triage without phenotyping; prioritizing isolates for AST; data-sharing advocacy

Within these limits, the results suggest a concrete operational role for context-only prediction: triage. Because the model requires no resistance-gene calls and no phenotyping, it can prioritize incoming isolates for confirmatory phenotypic AST — concentrating scarce laboratory capacity on the highest-risk contexts — and can flag high-risk lineages entering a country before their resistance cargo is characterized, provided the per-country transportability distribution (Section 3.1), not the pooled estimate, governs deployment decisions. The label-scarcity audit carries a broader policy implication: with only 2.43% of African KpSC records (and 1.1% of KpSC BioSamples globally) carrying submitter-supplied antibiograms, the surveillance community is sequencing far faster than it is phenotyping; routine antibiogram submission alongside genome deposition, through the existing NCBI BioSample antibiogram route^20^, would convert the genotypic-proxy workaround described here from a necessity into a choice. Finally, the pipeline itself — locked inclusion rules, a hard leakage gate at model entry, pinned and dated data releases, a frozen accession manifest, fixed seeds, and a 96-test suite with bit-for-bit reproducible reruns — is a reusable template for leakage-controlled AMR machine learning on drifting public archives, addressing the reproducibility deficiencies that reporting-guideline assessments have documented across this literature^35^.

## 4. Conclusion

### 4.1 Summary of Contributions

This study set out to determine whether the genomic *context* of African *Klebsiella pneumoniae* species complex (KpSC) isolates — lineage, capsule and O-locus background, and virulence loci, with every resistance determinant excluded — carries enough signal to predict aggregate multidrug resistance (MDR) status, and whether that signal survives geographic transport. Three contributions follow.

First, we constructed an AMR-gene-free MDR prediction framework for African KpSC, and we answered the two open questions that determine whether such a framework is even feasible on public data. Phenotypic linkage is extremely scarce: only 231 of 9,505 strict-*K. pneumoniae* African records (2.43%) carry submitter-supplied antibiograms, concentrated in six countries and seven BioProjects, which forces a rule-based genotypic MDR proxy as the primary label. Country sufficiency is nonetheless adequate for geographic validation: eight countries clear n ≥ 200 genomes and fourteen clear n ≥ 100, supporting a leave-one-country-out (LOCO) design on the current snapshot but not on any previously published cohort.

Second, we performed a LOCO cross-validation of AMR machine learning in *K. pneumoniae* or any Enterobacterales. The pilot demonstrates the expected random-to-LOCO generalization gap: pooled area under the receiver operating characteristic curve (AUROC) of approximately 0.85 under random splitting fell to 0.66–0.69 under LOCO across three model families, with wide per-country dispersion (0.55–1.00) and single-class folds reported as non-estimable rather than dropped. Pilot fold sizes (n ≤ 20 test genomes) make these point estimates illustrative; the substantive result is the ordering of schemes and the demonstration that the gap exists and is measurable.

Third, we delivered a leakage-controlled, reproducible pipeline — 96/96 passing tests, a pinned toolchain (Kleborate 3.1.3, Kaptive 3.0.0b6, mash 2.3, minimap2 2.31, Python 3.12), fixed seed 20260821, and a hard leakage guard at model entry — together with an independent audit of the descriptive baseline article (88,183 global and 3,796 African supplementary rows verified; a publisher PDF mis-bundling reported to the journal). The operative test for genomic AMR surveillance models, this study argues, is geographic validation, not pooled accuracy.

### 4.2 Future Work

The pilot establishes feasibility; the full study is the immediate next step. Specifically: (i) scaling to the full 3,000–5,000-genome cohort across the fourteen qualifying countries, which will narrow per-fold confidence intervals and make the per-country LOCO distribution a confirmatory rather than illustrative endpoint; (ii) adding Panaroo pangenome (gene presence/absence) features with explicit AMR decontamination — removal of all resistance-associated gene clusters — to test whether accessory-genome context adds signal beyond lineage and capsule; (iii) incorporating BIGSdb-Pasteur LIN-code features as a finer-grained, hierarchical lineage encoding; (iv) quantifying genotype–phenotype label agreement (Cohen’s kappa) on the 231-isolate AST subset to bound the measurement error of the genotypic proxy label; (v) prospective validation on newly deposited African genomes as a temporal hold-out complement to the geographic hold-out; and (vi) archiving the repository and frozen data snapshots under a Zenodo DOI upon public release.

## 5. Methods

The complete analytical pipeline is summarised in Figure 1. All design decisions described below were fixed before model fitting and are implemented in a versioned, test-covered code base (Section 5.7).

**Figure 1**. Overview of the data-acquisition, typing, labelling, feature-engineering, validation, and explainability pipeline. Predictor construction (genomic context) is deliberately separated from label construction (acquired-resistance evidence), and every feature matrix must pass an automated leakage assertion before entering any model.

Figure 1 illustrates the five stages of the study: (1) acquisition of three data sources with explicit versioning and integrity checks; (2) quality control (QC), species confirmation, and typing with a pinned toolchain; (3) construction of the MDR label under a strict acquired-resistance-only definition, alongside a leakage-controlled feature matrix; (4) a three-level validation hierarchy culminating in LOCO-CV; and (5) model training with pre-registered hyperparameters followed by block-level explainability with circularity controls. The diagram encodes the study’s central methodological commitment: because the label is itself derived from the genome, predictor construction is restricted to genomic *context* (lineage, capsular and lipopolysaccharide loci, virulence loci, plasmid replicons, decontaminated pangenome content, and metadata), while all direct resistance determinants and their aggregates are quarantined on the label side. The figure also records known data limitations (the publisher-side supplementary mis-bundle and the 2.43% AST coverage) that motivated downstream design choices such as the label hierarchy and the LOCO fold rules.

### 5.1 Data Sources and Cohort Definition

#### Reference supplements and audit findings

The primary sampling frame derives from the supplementary accession tables of the reference surveillance study by Ekwanzala^4^. We retrieved Supplementary Material 2 (MOESM2; XLSX) and Supplementary Material 3 (MOESM3; TSV) directly from the publisher’s supplementary listing and recomputed all headline counts locally: MOESM3 contains 88,183 data rows describing the global collection, and MOESM2 contains 3,796 African accessions spanning 22 countries^42^ ^18^ ^24^. Within MOESM2, Malawi (n = 1,247) and South Africa (n = 842) jointly account for 2,089 of 3,796 rows (55.0%), indicating pronounced country-level skew that any downstream validation design must accommodate^18^. Cross-referencing MOESM2 BioSample accessions against our independently downloaded National Center for Biotechnology Information (NCBI) metadata (Section 5.1) confirmed an overlap of 3,360 of 3,796 accessions (88.5%)^7^.

During the audit we identified a publisher-side packaging error: the body of the early-access PDF distributed under the reference article’s DOI and correct title page consists of an unrelated manuscript on geopolymer mortars, and the live article page exposes only the abstract^43^ ^39^ ^40^. Consequently, no Methods-level detail could be recovered from the reference article itself; we therefore treat MOESM2/MOESM3 as the authoritative record of that study’s accession set and disclose this limitation explicitly rather than inferring undocumented processing steps.

#### Current NCBI Pathogen Detection snapshot

The second data source is the NCBI Pathogen Detection Klebsiella taxon-group metadata export, pinned to release PDG000000012.2501^14^. The archived tab-separated value (TSV) file contains 172,412 Klebsiella-group records; filtering to strict *Klebsiella pneumoniae* with African country annotations yields 9,505 isolates, of which 231 (2.43%) carry a non-empty AST_phenotypes field^7^ ^14^. Because Pathogen Detection releases are regenerated continuously, the pinned release file was downloaded once, checksummed, and archived locally with the analysis code; a second, independent pull taken one day later showed approximately 3% record-count drift (e.g., South Africa 2,543 versus 2,465 isolates), confirming that release pinning—not the live portal—is the only reproducible basis for cohort definition^44^ ^41^. The AST_phenotypes strings originate from submitter-supplied BioSample antibiograms and are not vetted or breakpoint-harmonised by NCBI; interpretation therefore follows the label-hierarchy rules in Section 5.3^20^ ^45^ ^46^.

#### Pilot cohort

To validate the toolchain end to end before full-cohort processing, we constructed a stratified pilot cohort of 180 genomes: 20 genomes per country across nine African countries, sampled with the global random seed 20260821. Assemblies were retrieved from the European Nucleotide Archive (ENA) using run-level queries against the *K. pneumoniae* species-complex taxon set^47^, and every downloaded file was verified against its archive-supplied MD5 checksum before acceptance; failed downloads were re-fetched once and otherwise excluded. The pilot cohort is used for pipeline verification (QC gate behaviour, parser correctness against the Kleborate v3 output schema, label-construction unit tests) and is not pooled into the primary analysis.

We note explicitly that EnteroBase was evaluated as a candidate source and rejected: it hosts no Klebsiella database, so the cgMLST analogue for this species complex is the Institut Pasteur BIGSdb scheme^48^ ^49^. The AllTheBacteria resource, which provides 75,291 uniformly assembled *K. pneumoniae* genomes with precomputed Kleborate v2.3.2 typing, was likewise not mixed into the primary cohort, because its typing schema (Kleborate v2.3.2) is not interchangeable with the locally run Kleborate v3.1.3 output (Section 5.2)^50^ ^51^ ^52^. Table 1 summarises the data sources and their roles.

Table 1 makes explicit the asymmetry that shapes the entire design: accession-level metadata are abundant (hundreds of thousands of records), sequence data are retrievable at scale, but phenotypic AST coverage is rare (231 of 9,505 African strict-Kp records) and geographically concentrated. The reference supplements contribute a curated African sampling frame but arrive with a documented publisher-side integrity problem, so their role is restricted to accession enumeration and country composition. The pinned Pathogen Detection release is the authoritative cohort source, and the ENA pilot verifies that every software and parsing step functions on real assemblies of the target population before the full download is attempted. AllTheBacteria is retained as contextual evidence only; mixing its v2.3.2 typing calls with local v3.1.3 output would introduce an unquantified schema-confound into both labels (resistance-class counts) and features (K/O locus nomenclature), which the study’s leakage-control rules prohibit.

### 5.2 Species Confirmation, Typing, and QC

#### QC thresholds

Assemblies were required to pass three gates chosen to match established surveillance conventions for the *K. pneumoniae* species complex: total assembly length 4.5–7.0 Mb, at most 500 contigs, and N50 ≥ 20 kb. These bounds follow the QualiBact framework as deployed in Pathogenwatch, whose published KpSC thresholds (N50 ≥ 15 kb; 4.6–6.4 Mb; ≤500 contigs) we widened slightly on genome length to avoid excluding legitimately large, plasmid-rich MDR genomes—a population of direct interest here—while retaining a stricter N50 gate because fragmentation propagates into untypeable capsule-locus calls^53^ ^54^ ^55^. Assembly statistics (total length, contig count, N50, GC fraction) were computed with a built-in implementation equivalent to QUAST’s core metrics; a QUAST wrapper is available for optional confirmation^56^. Isolates failing any gate were excluded from feature construction but retained in a QC-fail log with reasons.

#### Pinned typing toolchain

Species confirmation, sequence type (ST), capsule (K) and lipopolysaccharide (O) locus typing, virulence-locus detection, and resistance-determinant calling were performed with Kleborate v3.1.3 in -p kpsc mode, which invokes Kaptive 3.0.0b6 for K/O locus assignment and mash 2.3 with minimap2 2.31 for species screening and alignment-based typing^23^ ^57^ ^58^ ^23^. The toolchain is pinned at the level of every dependency: Kaptive was fixed at 3.0.0b6 because kaptive ≥ 3.0.1 removed the kaptive.database application programming interface (API) that Kleborate 3.1.x imports, a version interaction we validated locally before locking the environment; this pin combination has independent precedent in recent African neonatal-unit genomic work^59^. Parsers target the Kleborate v3 output schema (file klebsiella_pneumo_complex_output.txt), whose renamed Kaptive columns and CARD v3.2.9-based resistance database differ from the v2 schema; mixing v2-era precomputed calls with local v3.1.3 output is therefore prohibited^58^ ^23^ ^52^. Kaptive v3 untypeable calls are treated as data, not errors, because assembly fragmentation is a recognised cause of untypeable K loci and such calls carry QC information (Section 5.4)^55^.

### 5.3 MDR Label Construction

#### Acquired-resistance-only definition

The prediction target is binary MDR status under the international expert definition of Magiorakos et al.: non-susceptibility to at least one agent in three or more antimicrobial categories^5^ ^5^. Critically, that definition applies to *acquired* resistance, and its correct application to *K. pneumoniae* requires excluding intrinsic determinants^60^ ^61^. Naive gene counting violates this clause: in our snapshot audit, counting all AMRFinderPlus/Kleborate resistance hits—including intrinsic fosA, oqxAB, emrD, and the core chromosomal blaSHV conferring intrinsic ampicillin resistance—flags 99.3% of the cohort as MDR, whereas counting only acquired classes yields 87.6%^33^ ^62^ ^63^. We therefore excluded fosA, oqxAB, emrD, and wild-type blaSHV from all class counts; the Kleborate resistance-class counter already implements curated exclusions of intrinsic ampicillin, chromosomal β-lactamase and porin-mutation artefacts, which is a further reason for preferring it as the genotypic source^63^ ^36^.

#### Label hierarchy and provenance

Labels follow a strict source hierarchy, recorded per isolate in a label_source provenance field (phenotypic_ast, genotypic_proxy, or excluded). Phenotypic AST labels are preferred whenever the AST_phenotypes string parses and the panel passes a completeness rule (minimum number of drugs and classes tested, recorded as n_drugs_tested/n_classes_tested; zero-valued number_drugs_* counters in the NCBI schema are missing-data sentinels, not pan-susceptibility). Because EUCAST’s 2019 redefinition renders “I” as “susceptible, increased exposure” whereas CLSI retains “intermediate”, and because submitter-supplied public AST strings cannot be retrospectively harmonised to a single breakpoint era, primary labels count resistant (R) calls only; an R+I non-susceptibility label is pre-registered as sensitivity analysis SA1^64^ ^65^ ^66^ ^67^. When AST is absent or uninterpretable, the fallback is the Kleborate genotypic proxy num_resistance_classes ≥ 3, which has published precedent for MDR assignment in *K. pneumoniae* genomic epidemiology^68^ ^57^. Isolates for which neither route yields a label are marked excluded and dropped from supervised analyses.

#### AST scarcity and its consequence

Only 231 of 9,505 (2.43%) African strict-Kp records carry any AST string, drawn from six countries and a small number of resistance-enriched BioProjects^7^ ^69^. This scarcity, combined with the enrichment of AST availability towards resistant and outbreak isolates, means phenotypic labels cannot support the primary analysis without severe selection bias; a systematic review of genotype-based AST prediction reaches the same conclusion regarding phenotype-label quality in public repositories^70^ ^46^. Consequently, AST-derived labels are reserved for (i) a phenotype-only sensitivity cohort and (ii) label-agreement (Cohen’s κ) checks against the genotypic proxy, while the primary cohort uses the genotypic proxy with the intrinsic-exclusion rules above. This is reported as a deliberate bias–coverage trade-off, not as equivalence between label sources; known genotype–phenotype discordance mechanisms (alternative resistance pathways, silent genes) remain a limitation ^38^ ^71^.

### 5.4 Feature Engineering and Leakage Control

#### Context feature blocks

Predictors comprise five genomic-context blocks plus QC/metadata: (i) ST/lineage indicators from the seven-locus Pasteur multilocus sequence typing (MLST) scheme (one-hot with minimum-count binning of rare STs into other)^72^ ^73^; (ii) K- and O-locus assignments, with untypeable calls retained as their own category because they are biologically and technically informative rather than missing^74^ ^75^ ^55^; (iii) virulence-locus features (yersiniabactin, aerobactin, salmochelin, colibactin, hypermucoidy loci) from Kleborate’s virulence modules, motivated by documented convergence of virulence and MDR plasmids in this species^76^ ^77^ ^78^ ^79^; (iv) plasmid-replicon features (Section 5.4); and (v) a pangenome presence/absence block built with Panaroo, which we chose over Roary because its graph-based correction reduces spurious cluster fragmentation and accessory-genome inflation that would otherwise corrupt feature interpretability^80^ ^81^ ^82^ ^83^. The pangenome matrix passes through a versioned AMR-decontamination step—every gene cluster annotated as a resistance determinant by AMRFinderPlus or Kleborate is removed before feature construction—because the raw presence/absence matrix contains the very genes that define the label^84^. A core-genome single-nucleotide polymorphism (SNP) phylogeny (Panaroo core alignment, variable sites extracted with SNP-sites, maximum-likelihood inference with IQ-TREE 2 and ModelFinder with UFBoot2 support) is used for lineage annotation and confounding diagnostics; recombination handling with Gubbins is restricted to the lineage level, per its documented scope, and is not applied to concatenated core-gene alignments^85^ ^86^ ^87^ ^88^ ^89^ ^90^.

#### Forbidden derivatives

Because the genotypic label is computed from resistance-gene content, any feature derived from that content would constitute label leakage; benchmark experience shows that models trained with such derivatives achieve meaningless near-perfect accuracy^32^. The pipeline therefore enforces an explicit deny-list—resistance-gene columns, resistance scores, num_resistance_classes, and all amr_/resistance_/bla_-prefixed columns—checked programmatically by assert_no_label_leakage at model entry, so that no model can be fitted on a matrix containing a label derivative. Target encoding of categorical features is forbidden outside fold-internal fitting, consistent with established evidence that out-of-fold leakage of target statistics inflates error estimates^91^ ^92^. Country is used only for grouping (LOCO) and sensitivity analyses, never as a default predictor, because region proxies entrench rather than remove geographic bias in AMR prediction^25^.

#### Plasmid-replicon residual circularity

Plasmid-replicon features (PlasmidFinder/MOB-suite conventions) occupy an ambiguous position: they are legitimate genomic context, yet replicon–cargo coupling (e.g., IncF–blaCTX-M-15, IncX3–OXA-181 associations) makes them mechanistically adjacent to the resistance label^93^ ^94^ ^31^ ^95^. The coupling is strong but non-deterministic—replicons without cognate resistance cargo occur in African lineages—so replicons are retained as predictors under three pre-registered controls: an ablation ladder comparing models with and without the replicon block, label-permutation nulls (Section 5.6), and explicit interpretive framing of any replicon-driven signal as cargo-correlated context rather than independent evidence ^79^ ^31^. Population-structure confounding between lineage and resistance, demonstrated at scale across five pathogens including *K. pneumoniae*, is the motivating threat model for both the feature restrictions here and the validation hierarchy in Section 5.5^11^ ^11^.

### 5.5 Validation Design

#### Three-split hierarchy

Model performance is estimated under three nested regimes with distinct inferential meanings, following recent formal analyses of what cross-validation estimates^27^ ^96^. A random 80/20 split (seed 20260821) estimates interpolation within the sampled population—the regime in which nearly all published bacterial AMR machine learning operates and in which performance is known to be optimistic ^11^ ^12^. A lineage-aware split, in which all genomes of an ST fall entirely in training or test partitions (grouped cross-validation), estimates generalisation to unseen clones, addressing the clade-held-out failure mode documented for population-structure-confounded predictors ^27^ ^11^. LOCO-CV, in which all isolates from one country form the held-out fold, estimates geographic transportability—the property that multi-country evaluations have repeatedly shown to fail when models trained in one region are applied to African data^10^ ^10^. The primary comparison is the Wilcoxon signed-rank test over per-country AUROC values, with a bootstrap-over-countries confidence interval that treats countries, not genomes, as the resampling units^97^ ^98^.

#### LOCO fold rules

A country qualifies as a primary LOCO fold at n ≥ 200 genomes, because at that size the binomial standard error at the least favourable prevalence (p = 0.5) bounds 95% confidence half-widths to approximately ±7 percentage points; the n ≥ 50 alternative was rejected as uninformative. Under the pinned snapshot this yields eight primary folds (South Africa, Malawi, Kenya, Tanzania, Ghana, Tunisia, Nigeria, Ethiopia); countries with n ≥ 100 are analysed as secondary folds with wider intervals reported. Two folds carry an explicit caveat: Malawi’s and Ethiopia’s top three BioProjects contribute 97.3% and 90.3% of their genomes respectively, so those folds are effectively leave-one-study-out and are flagged as such in all per-fold reporting^17 59^. Leave-one-country-out designs have precedent in large-scale AMR burden estimation, where they serve the same transportability-estimation purpose^1^. In the pilot cohort (20 genomes per country), the full-cohort sufficiency thresholds cannot be met, so pilot LOCO folds instead apply the pipeline’s minimum-viability rule (implemented as leave_one_country_out(min_train=50, min_test=20)): a country forms a pilot fold only if at least 20 labelled test genomes and at least 50 training genomes remain after species confirmation and label construction, which is why three pilot countries were non-estimable (Section 3.3). Pilot folds are therefore illustrative only and are labelled as such wherever they are reported.

#### Models and class imbalance handling

Three estimator families are compared: logistic regression (a deliberately interpretable baseline, in line with arguments that high-stakes problems should be anchored by transparent models)^99^, random forest, and LightGBM. Class imbalance is handled with inverse-frequency class weights (class_weight="balanced" for logistic regression; equivalent weighting for the tree models), not by resampling: synthetic oversampling risks train–test contamination and area-under-curve inflation when any resampling precedes partitioning, and grouped geographic folds make neighbourhood-based synthesis ill-defined^100^ ^101^. All hyperparameters were fixed before any model saw evaluation data (pre-registered in the project configuration and recorded verbatim in the machine-readable run manifest, Section 5.7) to avoid selection bias in error estimation^91^ ^102^.

#### Metrics

The primary metric is the area under the receiver operating characteristic curve (AUROC) with bootstrap confidence intervals. Because MDR prevalence varies substantially by country, the area under the precision–recall curve (AUPRC) is reported alongside the fold-specific prevalence baseline, precision–recall analysis being the more informative view under class imbalance^103^ ^104 28^. The Matthews correlation coefficient (MCC) is reported as the threshold-dependent summary of confusion-matrix performance, given its documented advantages over accuracy and F1 under imbalance^26^ ^105^. Calibration is assessed with calibration curves and intercept/slope, following the STRATOS calibration hierarchy^106^ ^107^. Secondary metric families (per-antibiotic-class analyses, sensitivity analyses such as SA1) are multiplicity-controlled with Benjamini–Hochberg false discovery rate (BH-FDR) correction^108^.

### 5.6 Explainability

#### Block-level SHAP with three-route concordance

Explanations are computed at the level of feature blocks, not individual one-hot columns. For tree models we use TreeSHAP with interventional feature perturbation (feature_perturbation="interventional"), which evaluates marginal contributions against a background distribution rather than conditioning on observed feature values, reducing extrapolation into impossible feature combinations^29^ ^109^ ^110^ ^111^. The background dataset is 200–500 rows drawn from the training portion of each fold only—fold-internal background—so that no information from held-out rows enters the explanation machinery; explanations are computed for held-out rows only. Block-level importances are triangulated across three routes: (i) TreeSHAP values summed within blocks; (ii) grouped permutation importance, in which an entire block is permuted jointly, computed on held-out data given the known failure modes of permutation on training data and under correlation^112^ ^113^ ^114^ ^115^; and (iii) the ablation ladder (Section 5.6). We report the caveat that summing one-hot SHAP values within a block is not, in general, equal to the exact grouped Shapley value, and treat concordance across the three routes—not any single method—as the evidence standard^116^ ^30^ ^117^.

#### Ablation ladder and label-permutation nulls

The ablation ladder refits each model while removing one feature block at a time (replicon block first, given its circularity risk), providing an explanation-by-removal estimate of each block’s contribution that is robust to the known failure modes of post hoc explainers^118^ ^119^. As circularity controls, label-permutation nulls (labels shuffled within fold structure, full pipeline refitted) are run pre-registered; any nominally significant result under permuted labels invalidates the corresponding real-label finding. Shortcut-learning precedents in clinical machine learning motivate treating these nulls as mandatory rather than optional^120^.

### 5.7 Reproducibility and Reporting

#### Reporting standards

The manuscript follows TRIPOD+AI, the applicable guideline for prediction-model studies using regression or machine learning; items specific to prospective or interventional designs are marked not applicable with justification, and the completed checklist accompanies the submission^121^ ^122^ ^123^. A PROBAST+AI self-assessment of risk of bias and applicability is provided as supplementary material^124^. The TRIPOD-Cluster extension informs our handling of country-clustered data^122^.

## Data and code availability

Reproducibility rests on five concrete artefacts: (i) accession-level supplements listing every genome used, satisfying mandatory individual-accession deposition norms^125^; (ii) the archived, checksummed Pathogen Detection release PDG000000012.2501 TSV, acknowledging measured inter-release drift^44^ ^41^; (iii) a pinned software environment (exact versions and database versions, per AMRFinderPlus-style guidance that both software and database versions be stated)^126^; (iv) a machine-readable run manifest recording the global seed 20260821 and all configuration values; and (v) a 96-test suite covering label construction, feature leakage, and split integrity. Code is deposited in a DOI-issuing repository archive at the release corresponding to this manuscript, because a version-control link alone does not meet publisher code-availability policy^127^ ^128^ ^129^. Data and code availability statements follow the drafted language aligned with TRIPOD+AI open-science items 18a–f^122^.

## Acknowledgements

The authors thank the contributors and curators of NCBI Pathogen Detection, NCBI BioSample, and BIGSdb-Pasteur, whose open data and typing schemes made this reanalysis possible, and the investigators of the primary African surveillance studies — particularly Ekwanzala and colleagues — for making their genome accessions and supplementary metadata publicly available. This work used the Kleborate, Kaptive, mash, and minimap2 software tools developed by their respective teams. During the preparation of this work, the authors used ChatGPT (OpenAI, GPT-5) and Claude (Anthropic, Claude Sonnet 4) to improve the readability and language clarity of the introduction and discussion sections. After using these tools, the authors reviewed and edited the content as needed and took full responsibility for the content of the publication. The authors also used Claude Opus 4.6 to assist in debugging and optimizing the Python script used for preliminary data cleaning. All outputs were independently verified. This research received no specific grant from any funding agency in the public, commercial, or not-for-profit sectors.

## Author contributions

Conceptualization: [A.A.]; Methodology: [all authors]; Software: [A.A.]; Validation: [E.B.]; Formal analysis: [A.A]; Investigation: [all authors; Data curation: [all authors]; Writing — original draft: [E.B.]; Writing — review & editing: [all authors]; Visualization: [A.A.]; All authors read and approved the final manuscript.

## Additional information

Correspondence and requests for materials should be addressed to the corresponding author.

## Data Availability

This study reanalyses publicly available genome assemblies and metadata; no new sequencing was performed. The primary data source is the NCBI Pathogen Detection *Klebsiella* taxon group release PDG000000012.2501 (metadata TSV downloaded 2026-08-20; https://ftp.ncbi.nlm.nih.gov/pathogen/Results/Klebsiella/); the exact filtered metadata snapshot used for all analyses is archived with this paper because public archives are continuously updated — we observed approximately 3% turnover in the largest national subset between two pulls of the same source^44^ ^41^. Reference accession tables (supplementary files MOESM2 and MOESM3) from Ekwanzala et al. (2026; doi:10.1038/s41598-026-60706-4) were used as an additional input and are cited as a data source in accordance with third-party data fair-use norms^125^. The pilot accession list (180 assembly accessions with BioSample and BioProject linkage, country, and label provenance) is provided as a supplementary table, in line with the requirement that individual accessions rather than umbrella BioProjects be listed^125^. Phenotypic antimicrobial-susceptibility data are those supplied by the original submitters to NCBI BioSample antibiogram fields and were not independently verified; these 231 records are used only for label-agreement analysis, not as the primary label source. All results correspond to the frozen, dated snapshots recorded in results/run_manifest.json.

## Code Availability

The complete analysis pipeline, klebsiella_amr_pipeline v0.1.0 — including data download, quality control, typing, label construction, feature engineering, split generation, model training, explainability, and figure code — is openly available at https://github.com/Ultramedico/klebsiella-amr-pipeline (release v0.1.0), with the frozen, validated runtime environment archived as release asset runtime_snapshot.tar.gz (MD5 928f38e2ee5d73c854563455ea2c5d02);, since a repository link alone does not provide a permanent identifier^127^ ^128^. The repository includes the design specification (SPEC.md), a 96-test unit and end-to-end suite that runs without network access, and a pinned environment (Kleborate 3.1.3, Kaptive 3.0.0b6, mash 2.3, minimap2 2.31, Python 3.12, LightGBM, SHAP) with per-run software and database versions recorded in results/run_manifest.json^130^. All stochastic components use random seed 20260821; deterministic reruns reproduce the reported metrics exactly.

